# Cross-Modal Training Using Xenium Spatial Transcriptomics Enables DINO-DETR Based Detection of Vascular Niches in H&E Whole-Slide Images

**DOI:** 10.64898/2026.03.17.712266

**Authors:** S Pranali, Ramya Alugam, Shashank Gupta, Nameeta Shah, Megha S. Uppin

**Affiliations:** Amaranth Medical Analytics, Bengaluru, India; Department of Pathology, Nizam Institute of Medical Sciences, Hyderabad, Telangana, India

## Abstract

**Background:** Tumor vasculature is a key driver of glioma progression, yet routine quantification depends on subjective histopathologic assessment or resource-intensive ancillary immunohistochemistry. A scalable, objective method for vascular phenotyping from routine histology remains an unmet need.

**Methods:** We leveraged 10x Genomics Xenium spatial transcriptomics data from a glioblastoma specimen to generate molecularly resolved annotations of GBM-associated endothelial cells and pericytes across 809,041 cells. These annotations were transferred to matched H&E-stained sections to train a DINO-DETR-based object detection model using a binary classification scheme (vascular vs. other). The model was validated on four independent Xenium patient slides and applied to a retrospective cohort of 119 diffuse gliomas spanning WHO grades 2-4 (oligodendroglioma, astrocytoma, and glioblastoma) with linked survival data.

**Results:** Binary vascular cell detection achieved a precision of 0.78, a recall of 0.63, and an F1 score of 0.70, with an overall accuracy of 98.6%. Orthogonal spatial validation confirmed that predicted vascular cells were preferentially localized within annotated blood vessel regions. In subtype-stratified survival analysis, high AI-derived vascular cell proportion was significantly associated with worse overall survival in astrocytoma patients (log-rank p < 0.019).

**Conclusion:** Cross-modal AI training using spatial transcriptomics enables scalable, molecularly informed vascular quantification directly from routine H&E slides. Within the astrocytoma subtype, where tumor grade is most heterogeneous and vascular phenotype most variable, objective vascular quantification provides independent prognostic information demonstrating the potential of spatially supervised deep learning to extract clinically meaningful microenvironmental signals from universally available histologic material.

## Introduction

Microvascular proliferation is among the most powerful histologic predictors of outcome in diffuse gliomas, yet its routine assessment remains subjective, semi-quantitative, and irreproducible across pathologists^1,25^. Glioblastoma (GBM), the most aggressive primary brain tumor with a median survival of approximately 15 months, is characterized by florid microvascular proliferation, a defining histologic feature that reflects hypoxia-driven neovascularization and is required for histologic diagnosis under the 2021 WHO classification of CNS tumors^2,26^. Beyond GBM, the vascular microenvironment varies meaningfully across the diffuse glioma spectrum: lower-grade astrocytomas and oligodendrogliomas exhibit distinct angiogenic phenotypes whose prognostic significance remains incompletely characterized^3,8^. Tumor vasculature is not merely a passive bystander; it actively sustains glioma progression by supplying oxygen and nutrients, providing a perivascular niche for stem-like tumor cells, modulating immune infiltration, and serving as the principal target of antiangiogenic therapies such as bevacizumab^4,6,5^. Despite this central role, routine quantification of the vascular compartment relies on histopathologic assessment of microvessel density and microvascular proliferation, evaluations that are inherently observer-dependent and difficult to standardize across institutions^1^. Ancillary immunohistochemistry (IHC) with endothelial markers such as CD31, CD34, or von Willebrand factor has been used to improve objectivity, but this approach carries significant practical limitations^7^. Single-marker IHC cannot distinguish endothelial cells from the pericytes and perivascular fibroblasts that constitute the full neurovascular unit, conflating biologically distinct populations with potentially different prognostic implications^6^. Multiplexed IHC or immunofluorescence panels can resolve these populations but are costly, technically demanding, and not routinely performed on archival material in clinical practice^9^. As a result, large retrospective cohort studies of vascular phenotyping in glioma have been methodologically constrained, limiting the prognostic and biologic insight that can be extracted from existing tissue archives^3,8^.

Spatial transcriptomics has emerged as a powerful approach for molecularly resolved tissue analysis, enabling simultaneous quantification of hundreds to thousands of gene transcripts at subcellular resolution within intact tissue sections^10,11^. In-situ platforms such as 10x Genomics Xenium assign spatially registered transcript counts to individual cells, enabling unambiguous annotation of cell types, including endothelial cells, pericytes, and perivascular fibroblasts, based on their molecular profiles rather than morphologic appearance alone^8,13^. This molecular precision makes spatial transcriptomics an ideal source of ground truth annotations for supervising computational models^14^. However, spatial transcriptomics is resource-intensive, requires specialized instrumentation, and cannot realistically be applied at the scale of routine clinical archives comprising hundreds or thousands of cases^10^. A critical translational gap therefore exists between the molecular resolution achievable with spatial transcriptomics and the practical scalability of hematoxylin and eosin (H&E)-stained whole-slide images, which are universally available for archival specimens.

Computational pathology offers a means of bridging this gap. Deep learning models trained on H&E images have demonstrated substantial capacity for cell detection, tissue segmentation, and outcome prediction across multiple tumor types, including gliomas^18,19,20^. Crucially, recent work has shown that molecular annotations derived from spatial transcriptomics platforms can be transferred to matched H&E images to supervise AI models, a cross-modal training strategy that encodes molecular specificity into a modality that is universally available at scale^14,15,12^. Transformer-based object detection architectures, in particular, have shown strong performance for cellular detection tasks in pathology, offering precise localization of individual cells within complex tissue backgrounds^16^. Whether this strategy can be applied to the neurovascular unit in glioma, enabling scalable, molecularly informed vascular quantification from routine H&E slides, has not been established.

In this study, we leveraged 10x Xenium spatial transcriptomics data from a GBM specimen to generate molecularly resolved annotations of endothelial cells and pericytes, and transferred these annotations to matched H&E-stained sections to train a DINO-DETR-based object detection model. A key objective was to demonstrate that molecularly resolved annotations derived from a small set of WSIs can be sufficient to train a performant detection model when spatial transcriptomics data provides high-fidelity, cell-level ground truth, circumventing the need for large manually annotated training sets that are impractical to generate for rare cell populations such as vascular cells. We validated the model on four independent Xenium patient slides and subsequently applied it to a retrospective cohort of 119 diffuse gliomas spanning the full WHO grade spectrum. Using AI-derived vascular cell proportions, we tested the hypothesis that objective, H&E-based vascular quantification provides prognostic information across diffuse glioma subtypes. Our findings demonstrate that cross-modal AI training using spatial transcriptomics enables scalable vascular phenotyping from routine histology and reveals a prognostically significant vascular signature in astrocytoma that is independent of standard clinical covariates.

## Methods

### Slide Digitization and Whole Slide Imaging

Formalin-fixed paraffin-embedded (FFPE) tissue sections from patients diagnosed with glioma were Containing H&E stained slides of brain tumours were retrospectively compiled over the period 2019–2023 at Nizam’s Institute of Medical Sciences (NIMS), Hyderabad. Those slides followed standard histopathological protocols.

The stained glass slides were digitized using high-resolution whole slide scanners, including Optimus 6X and Panoramic-250, to generate Whole Slide Imaging (WSI) images.

A total of 119 whole slide images were included in the study. These consisted of 61 glioblastoma cases, 36 astrocytoma cases, and 22 oligodendroglioma cases. All digitized slides were reviewed to ensure image quality and diagnostic accuracy prior to further analysis.

### Dataset Variables Description

The dataset included demographic, clinical, radiological, and pathological variables for each patient. Each case was assigned an anonymized case number to ensure patient confidentiality. Demographic information included patient age at diagnosis and sex. Clinical features (C/F) captured the presenting symptoms or clinical characteristics, while radiological findings summarized key imaging observations.

Pathological evaluation included the final diagnosis and tumour grading according to the World Health Organization (WHO) classification of central nervous system tumours. The anatomical site of the tumour and the glioma subtype (oligodendroglia, astrocytoma, or glioblastoma) were also recorded.

Immunohistochemically and molecular markers were included as part of the pathological assessment. These comprised the Ki-67 proliferation index expressed as a percentage of positively stained tumour cells, the presence of IDH1 R132H mutation status, ATRX protein expression status, and P53 mutation status. These markers were recorded as categorical variables indicating either positive/negative status or retained/loss of expression, as applicable. In addition, overall survival time was recorded as the duration from the date of diagnosis to the date of death or last follow-up, measured in months. Survival status indicated the patient’s outcome at the last follow-up, which was used for survival analysis and predictive modelling.

### Spatial Transcriptomics Data and Cell Annotation

A publicly available FFPE human glioblastoma (GBM) tissue section profiled on the 10x Genomics Xenium in situ platform was used as the primary training dataset^17^. The tissue was obtained from Discovery Life Sciences and processed using the Xenium Human Immuno-Oncology Profiling Panel (380 genes), with cell segmentation staining performed per the manufacturer’s protocol. Xenium Onboard Analysis version 2.0.0 detected 816,769 cells, with a median of 157 transcripts per cell across a tissue area of approximately 104.98 mm^2^. Graph-based cell clusters were generated by the Xenium platform and provided as part of the standard analysis output; no independent clustering was performed. Clusters were annotated by examining differentially enriched genes within each cluster against known cell type marker profiles. Clusters 15 and 17 were identified as vascular populations based on enrichment of GBM-associated endothelial and pericyte marker genes, respectively, and were assigned a unified “Vascular” label. All remaining clusters were designated as “Other.” This binary annotation scheme, Vascular versus Other, served as the ground-truth label set for training the cell type detection model.

### Region of Interest Selection and Patch Extraction

Seven regions of interest (ROIs) were manually selected from the Xenium tissue section to capture the spatial and cellular heterogeneity of the tumor microenvironment. ROI areas ranged from 1.06 to 4.87 mm^2^. From these ROIs, image patches of 480 × 480 pixels were extracted, yielding a total of 4,712 patches for model training. Across all patches, 349,797 annotated cells were represented: 9,597 vascular cells (gbmEndo n = 6,351; gbmPeri n = 3,246; 2.7% of total) and 340,200 non-vascular cells (97.3% of total), reflecting the inherent rarity of vascular populations within the GBM tumor microenvironment and the class imbalance inherent to this detection task.

### Cell Type Detection Model Training

A DINO-DETR–based object detection model was trained to detect and classify vascular versus non-vascular cells within H&E-stained tissue patches. DINO-DETR (Detection with Transformers) is a transformer-based architecture that employs deformable attention and contrastive denoising training to achieve precise object localization in complex, densely populated scenes — properties well-suited to cellular detection in histopathology images^21^. The DINO_4scale_swin configuration was used, in which a Swin Transformer backbone extracts features across four spatial scales, enabling simultaneous capture of fine-grained cellular morphology and broader tissue context. Spatial transcriptomics annotations were transferred to matched H&E-stained sections to provide cross-modal supervision, encoding molecular cell type specificity into a universally available imaging modality. The model was trained on patches derived from a single WSI using an 80/20 train–test split over 50 epochs with a batch size of 4; training and validation loss both converged by approximately epoch 20–25, confirming that 50 epochs were sufficient for stable model convergence. H&E-specific data augmentation transforms were applied during training to improve generalization to unseen slides. Transfer learning was not employed. The best-performing model checkpoint was selected at epoch 48 based on validation loss.

For inference, input patches were resized to 480 pixels with a padding size of 32, and tiles of 416 pixels were processed with a batch size of 2. A confidence threshold of 0.2 was applied for detection, and an overlap threshold of 0.2 was used for non-maximum suppression. Class-specific inflection points were computed but not applied during inference. The final model configuration used the DINO_4scale_swin architecture.

### Model Validation on Held-Out Test Set

Model performance was first evaluated on the held-out 20% test set derived from the training WSI. Detection performance was assessed using standard object detection metrics: precision, recall, F1 score, and mean average precision (mAP) computed at an IoU threshold of 0.5. Given the binary classification scheme, metrics were calculated separately for the Vascular and Other classes. Overall pixel-level accuracy was computed as the proportion of correctly classified detections across both classes.

### Orthogonal validation on Xenium data

For orthogonal validation on independent patient data, Xenium WSIs from^22^ were used, which provide spatially resolved cell type annotations for human GBM tissue at subcellular resolution. The AI-predicted vascular cells from our model were mapped onto these slides, and a permutation-based spatial neighborhood analysis was performed to assess whether predicted vascular cells were preferentially co-localized with molecularly annotated vascular cells from the Xenium data. For each predicted vascular cell, the nearest Xenium-annotated cell type within a 100-pixel radius was identified. To establish a null distribution, the Xenium cell type labels were randomly shuffled 100 times, and the neighborhood analysis was repeated at each iteration. The observed distribution of nearest-neighbor cell type assignments, specifically the enrichment of Xenium-annotated vascular cells in the neighborhood of AI-predicted vascular cells, was then compared against the shuffled null distribution. Significant enrichment above the null would indicate that the model has learned spatially coherent vascular identity rather than detecting morphologically similar non-vascular cells by chance.

### Application to a Retrospective Glioma Cohort

The trained model was applied to H&E-stained WSIs from a retrospective cohort of 119 patients with diffuse glioma diagnoses spanning the full WHO grade spectrum, including glioblastoma (WHO grade 4), astrocytoma, and oligodendroglioma. Following inference, a sanity check was applied: in cases where multiple cell types were assigned to the same spatial location, the cell type with the highest predicted probability was retained as the final assignment. Cell type proportions per sample were then calculated as the fraction of each detected cell type relative to the total number of detected cells in the slide.

### Vascular Cell Type Analysis and Statistical Methods

The proportion of vascular cells per sample was calculated as the fraction of vascular detections relative to the total number of detected cells in that slide. Vascular proportions were compared across clinicopathologic and molecular subgroups, including glioma histologic subtype, WHO grade, IDH mutation status, ATRX mutation status, and p53 expression status. For comparisons across more than two groups, one-way ANOVA was used; pairwise comparisons between two groups were performed using an unpaired two-sided t-test.

For survival analysis, vascular proportion was dichotomized at the cohort median into high and low groups. Kaplan–Meier survival curves were generated for each diagnostic subtype independently (glioblastoma, astrocytoma, oligodendroglioma) and differences between strata were assessed using the log-rank test. Subtype-stratified analyses were performed to avoid confounding by the strong association between histologic subtype and both vascular activity and survival. Multivariate Cox proportional hazards regression was performed within the astrocytoma subgroup, with vascular class (high vs. low), age, and sex included as covariates, to determine whether AI-derived vascular proportion provided prognostic information independent of standard clinical variables. A second Cox model was fitted in the full cohort with WHO grade included as an additional covariate to assess the relationship between vascular proportion and grade as competing prognostic variables. Model discrimination was evaluated using the concordance index. All statistical analyses were performed in R.

## Results

To enable scalable, molecularly informed vascular quantification from routine H&E-stained whole slide images, we developed a cross-modal training framework in which spatial transcriptomics annotations from a single glioblastoma specimen were used to supervise a DINO-DETR-based object detection model (Figure 1). We first describe the generation of training data and model performance, followed by orthogonal validation in independent Xenium datasets, and finally, the application of the model to a retrospective cohort of 119 diffuse gliomas with linked survival data.

**Figure 1:**
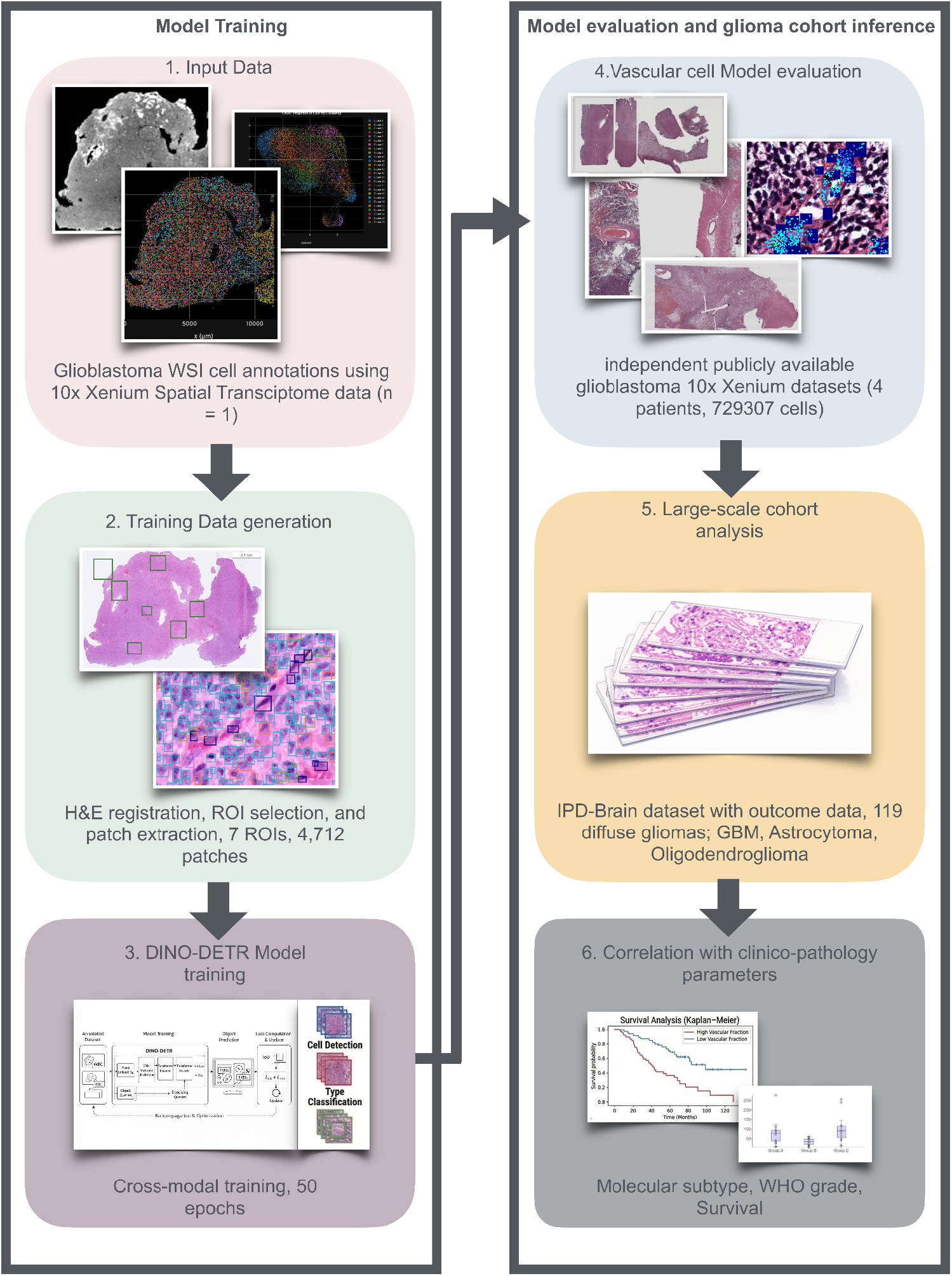
Overview of the study design.

### Spatial Transcriptomics-Guided Cell Annotation

Graph-based clustering of Xenium spatial transcriptomics data from a GBM tissue section yielded 26 spatially coherent cell clusters across 809,041 annotated cells (Figure 2A, 2B). To identify vascular populations, clusters were interrogated for differential enrichment of established vascular marker genes. Clusters K15 and K17 showed selective enrichment of endothelial markers VWF and CAV1, and pericyte/smooth muscle markers PDGFRB, ACTA2, and MYH11, respectively, and were jointly assigned a Vascular label (Figure 2C). All remaining clusters were designated Other. UMAP visualization confirmed that the Vascular-labeled cells occupied a distinct transcriptional neighborhood, segregating clearly from the broader tumor and immune cell populations (Figure 2D). Across all 809,041 annotated cells, vascular cells (gbmEndo n = 6,351; gbmPeri n = 3,246) comprised approximately 1.2% of the total tissue cellularity (Figure 2E), consistent with their known spatial restriction to vessel walls within the glioblastoma microenvironment. This pronounced class imbalance, with vascular cells representing a small minority against a dominant non-vascular background, defined the primary challenge of the subsequent detection task.

**Figure 2:**
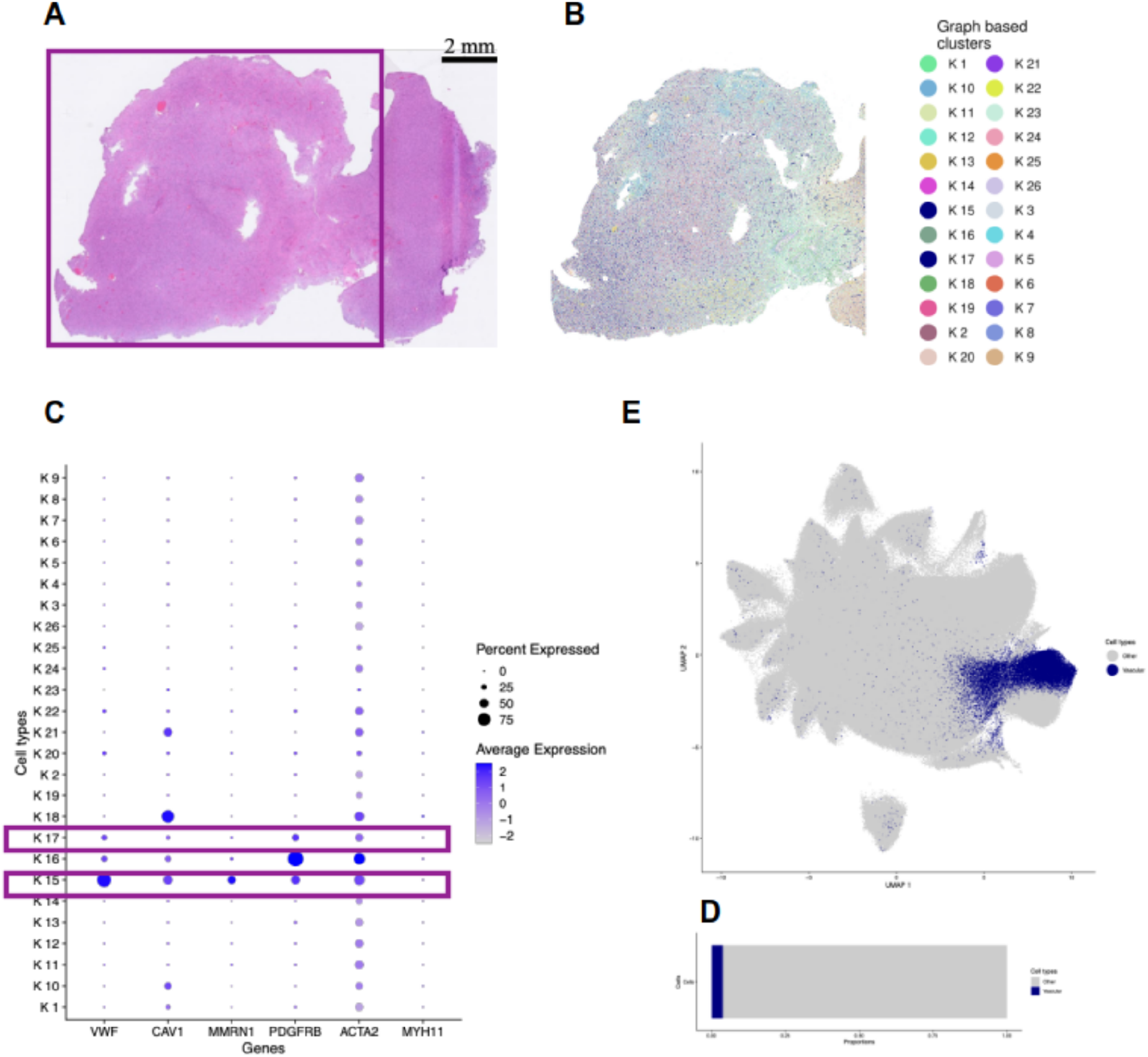
Xenium spatial transcriptomics-guided annotation of vascular cell populations in a GBM tissue section. (A) H&E-stained WSI of the GBM tissue section used for model training, shown at 2 mm scale. The tissue area profiled by the Xenium platform is indicated by the pink bounding box. (B) Spatial overlay of graph-based cluster assignments (K1-K26) on the Xenium tissue section, demonstrating spatially coherent cluster distributions reflecting the underlying tissue architecture. (C) UMAP projection of all 809,041 annotated cells, with Vascular cells (clusters K15 and K17, navy) highlighted against the Other population (grey). Vascular cells occupy a discrete transcriptional neighborhood, segregating clearly from tumor and immune populations. (D) Proportional bar chart showing the fraction of Vascular versus Other cells across the full tissue section; vascular cells comprise approximately 1.2% of total annotated cellularity. (E) Dot plot showing average expression and percent expression of canonical vascular marker genes, VWF, CAV1, MMRN1, PDGFRB, and ACTA2, and MYH11, across all 26 graph-based clusters. Clusters K15 and K17, highlighted by maroon boxes, show selective enrichment of endothelial and pericyte marker genes, respectively, supporting their annotation as GBM-associated vascular populations.

### Model Training and Performance

Seven ROIs were manually selected across the GBM tissue section to capture the spatial and cellular heterogeneity of the tumor microenvironment (Figure 3A). A representative high-magnification view of a training patch illustrates the overlay of ground truth annotations, with vascular cells marked by dark blue bounding boxes and non-vascular cells by grey-green boxes (Figure 3B). The DINO-DETR model converged smoothly over 50 epochs, with both training and validation loss declining steeply in early epochs and plateauing by approximately epoch 20–25 (Figure 3C, left). Validation loss was consistently lower than training loss throughout training, reflecting the application of H&E-specific data augmentation, which increases task difficulty relative to unaugmented inference patches. Mean average precision (mAP) increased steadily over training, reaching a plateau consistent with model convergence, with the best checkpoint selected at epoch 48 (Figure 3C, right). On the held-out 20% test set, binary vascular detection achieved a precision of 0.78, a recall of 0.63, and an F1 score of 0.70, with the Other class achieving near-perfect precision and recall (Figure 3D, left). The confusion matrix confirmed that of 1,967 true vascular cells, 1,247 were correctly detected, while 720 were missed and 351 non-vascular cells were incorrectly classified as vascular (Figure 3D, right). The lower recall relative to precision reflects the inherent challenge of detecting a rare cell population in a highly class-imbalanced tissue context.

**Figure 3.**
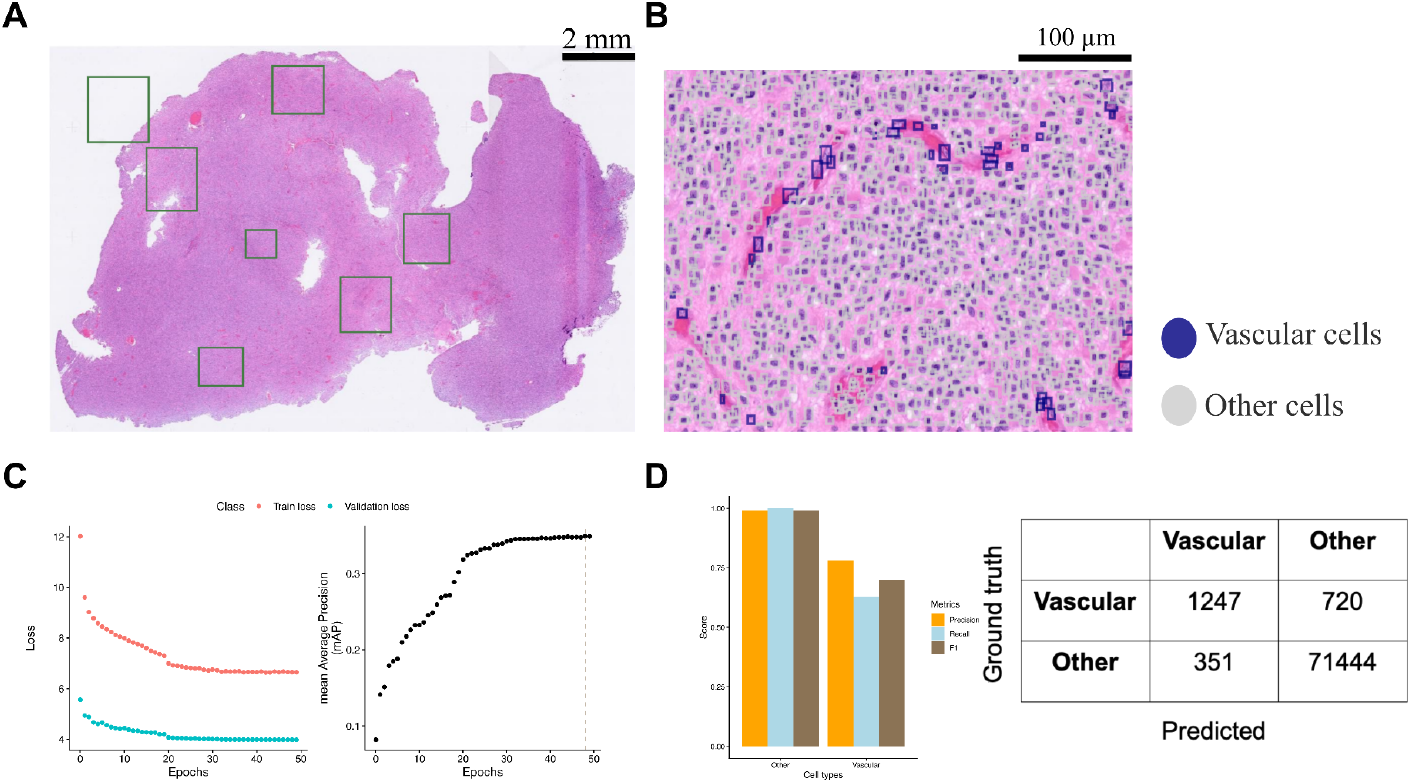
DINO-DETR model training, convergence, and binary vascular detection performance. (A) H&E-stained whole slide image of the GBM training section at 2 mm scale, with seven manually selected regions of interest (ROIs) indicated by green bounding boxes. (B) Representative high-magnification view (100 µm scale) of an H&E training patch with ground truth annotations overlaid; dark blue boxes indicate vascular cells and grey-green boxes indicate non-vascular cells. (C) Training and validation loss curves over 50 epochs (left); both curves converge by approximately epoch 20–25, with validation loss consistently below training loss, reflecting the effect of H&E-specific data augmentation applied during training. Mean average precision (mAP) over 50 epochs (right); mAP increases steadily and plateaus, consistent with model convergence, with the best checkpoint selected at epoch 48 (dashed vertical line). (D) Per-class precision, recall, and F1 scores for the Vascular and Other classes on the held-out test set (left); the Other class achieves near-perfect scores while the Vascular class reflects the challenge of rare-cell detection in a class-imbalanced setting. Confusion matrix showing raw detection counts for Vascular and Other classes (right); of 1,967 true vascular cells, 1,247 were correctly detected (true positives), 720 were missed (false negatives), and 351 non-vascular cells were incorrectly classified as vascular (false positives).

### Orthogonal Spatial Validation

To independently confirm that the model was detecting biologically meaningful vascular structures rather than morphologically similar non-vascular cells, the trained model was applied to three independent GBM Xenium WSIs (four patients) from^22^. (Nature Communications, 2026), which provide spatially resolved molecularly annotated cell type labels at the subcellular resolution. AI-predicted vascular cells were overlaid onto the Xenium slides, demonstrating widespread detection of vascular cells across all four patient specimens with spatial distributions consistent with known vessel architecture (Figure 4A, 4B, 4C). To quantitatively assess spatial concordance, a permutation-based neighborhood analysis was performed. For each AI-predicted vascular cell, the nearest Xenium-annotated endothelial or pericyte cell within a 100-pixel radius was identified. The observed distances between AI-predicted vascular cells and Xenium-annotated vascular cells were markedly shorter than those generated by randomly shuffling Xenium cell type labels 100 times, confirming significant spatial enrichment above the null distribution (Figure 4D). A representative high-magnification overlay further confirmed that AI-predicted vascular detections were spatially coincident with cells expressing canonical endothelial marker transcripts, including CD93, ESM1, PECAM1, and VWF (Figure 4E). Collectively, these results confirm that the cross-modal training strategy successfully encoded spatially coherent vascular identity into the H&E-based detection model.

**Figure 4.**
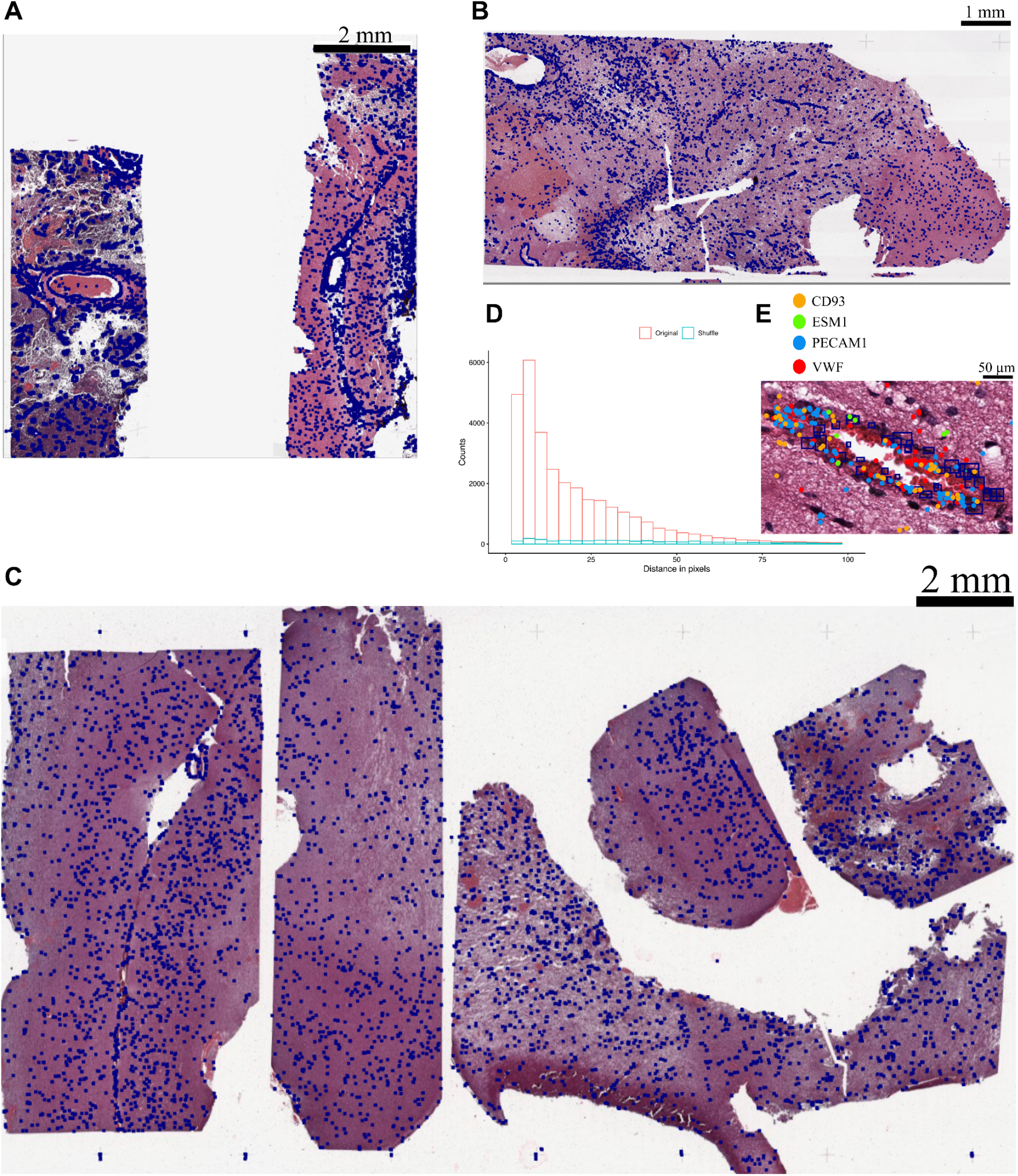
Orthogonal spatial validation of AI-predicted vascular cells in independent GBM Xenium datasets. (A, B, C) Representative Xenium whole slide images from four independent GBM patients^22^ ( Nature Communications, 2026) with AI-predicted vascular cells overlaid as dark blue dots, shown at 2 mm and 1 mm scale, respectively. (D) Permutation-based neighborhood analysis showing the distribution of distances between AI-predicted vascular cells and the nearest Xenium-annotated endothelial or pericyte cell in the original data (pink) versus 100 label-shuffled null permutations (teal). The pronounced left shift of the observed distribution relative to the null confirms that AI-predicted vascular cells are significantly enriched in the spatial neighborhood of molecularly annotated vascular cells. (E) High-magnification spatial overlay (50 µm scale) showing AI-predicted vascular bounding boxes co-localizing with cells expressing canonical endothelial marker transcripts CD93, ESM1, PECAM1, and VWF, confirming that model detections correspond to molecularly verified endothelial cells.

### AI-Derived Vascular Proportions Across the Glioma Spectrum

The validated model was applied to 119 diffuse glioma WSIs spanning WHO grades 2-4 (oligodendroglioma n = 21, astrocytoma n = 35, glioblastoma n = 63)^24^. Representative H&E patches from each diagnostic subtype illustrate the range of vascular cell densities captured by the model, with orange dots marking AI-predicted vascular cells overlaid on the tissue; insets indicate the location of each zoomed region within the full WSI (Figure 5A, 5B, 5C). Vascular cell proportions varied significantly across histologic subtypes (ANOVA p = 1.1×10^−8^), with glioblastoma showing the highest vascular proportions and oligodendroglioma the lowest, consistent with the known association between tumor grade and angiogenic activity (Figure 5D). Vascular proportions also differed significantly by WHO grade (ANOVA p = 1.3×10^−2^) and IDH mutation status (t-test p = 1.1×10^−7^), with IDH-wildtype tumors showing higher vascular proportions than IDH-mutant tumors. A significant difference was also observed by ATRX status (t-test p = 0.025). No significant association was observed with p53 expression status. Kaplan–Meier survival analysis stratified by vascular proportion within each diagnostic subtype demonstrated divergent survival trajectories, with the most pronounced separation observed in the astrocytoma subgroup (Figure 5E).

**Figure 5.**
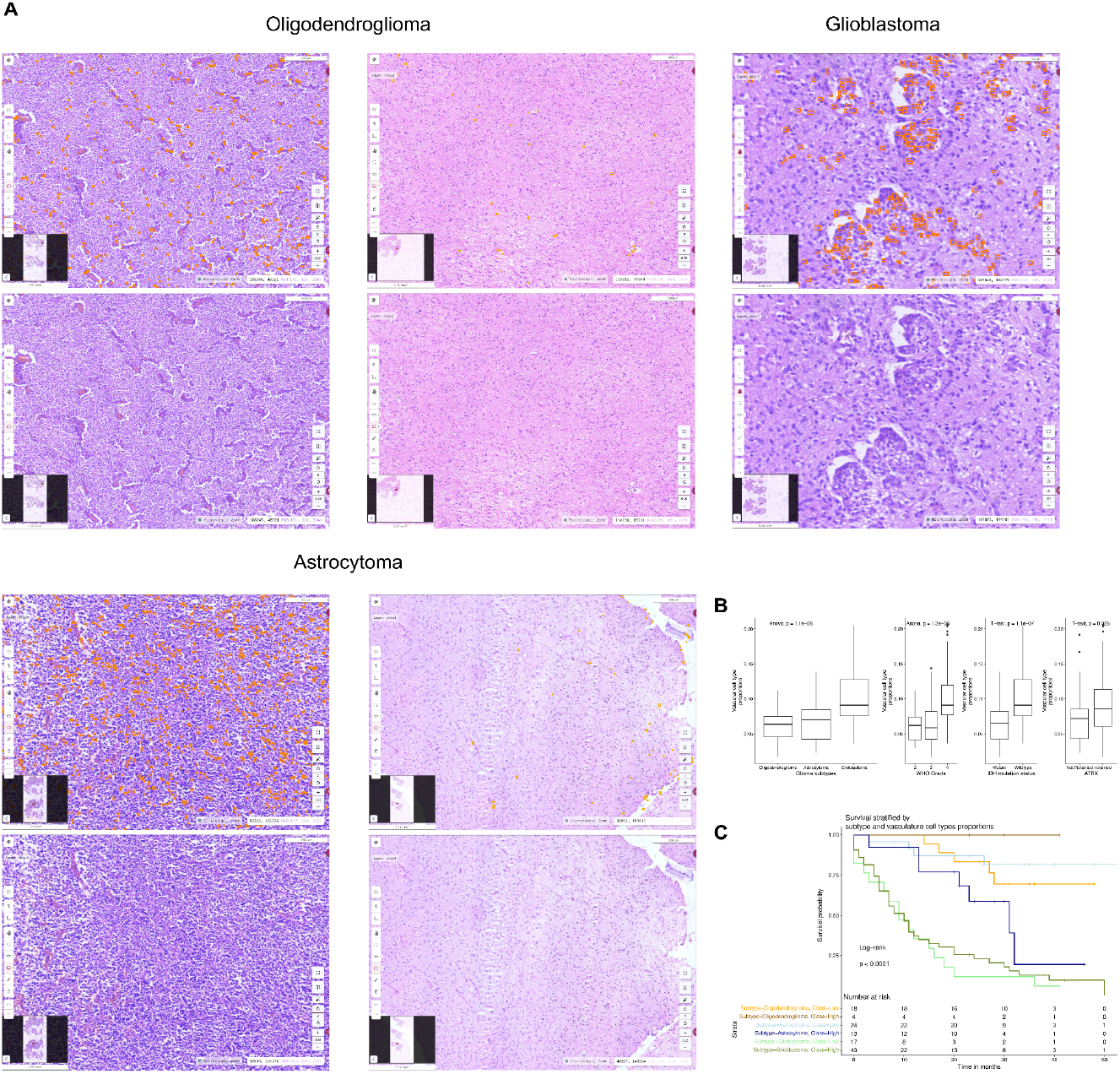
AI-derived vascular cell proportions across the diffuse glioma spectrum and association with survival. (A) Representative H&E patches from oligodendroglioma cases showing a vascular-high region (top left, with orange vascular cell overlay; bottom left, without overlay) and a vascular-low region (top right, with overlay; bottom right, without overlay). Insets indicate the location of each zoomed region within the full WSI. (B) Representative H&E patches from glioblastoma cases with (top) and without (bottom) vascular cell overlay, illustrating the dense vascular cell detection characteristic of GBM. The inset indicates the WSI location. (C) Representative H&E patches from astrocytoma cases showing a vascular-high region (top left, with overlay; bottom left, without overlay). The inset indicates the WSI location. (D) Box plots showing the distribution of AI-derived vascular cell proportions stratified by glioma histologic subtype (ANOVA p = 1.1×10^−8^), WHO grade (ANOVA p = 1.3×10^−2^), IDH mutation status (t-test p = 1.1×10^−7^), and ATRX mutation status (t-test p = 0.025). (E) Kaplan–Meier survival curves for all six subtype-vascular class strata (oligodendroglioma low/high, astrocytoma low/high, glioblastoma low/high), illustrating divergent survival trajectories across diagnostic subtypes stratified by AI-derived vascular proportion.

Collectively, these results demonstrate that cross-modal training using spatial transcriptomics annotations from a single glioblastoma specimen is sufficient to train a performant H&E-based vascular cell detection model. The model generalizes to independent patient slides, detects biologically coherent vascular structures as confirmed by orthogonal molecular validation, and produces vascular proportion estimates that recapitulate known associations with glioma grade and molecular subtype at the cohort scale. Critically, AI-derived vascular proportion provides independent prognostic stratification within the astrocytoma subtype, the subgroup where tumor grade is most heterogeneous and where objective vascular quantification adds clinically meaningful information beyond standard histologic classification.

## Discussion

In this study, we demonstrate that cross-modal training using spatial transcriptomics enables a DINO-DETR-based object detection model to identify vascular cell populations, specifically GBM-associated endothelial cells and pericytes, directly from routine H&E-stained WSIs. Applied to a retrospective cohort of 119 diffuse gliomas, AI-derived vascular cell proportions provided independent prognostic stratification within the astrocytoma subtype, identifying a high-vascular subgroup with significantly worse overall survival that was not explained by age or sex alone. These findings establish a proof-of-concept for spatially supervised computational vascular phenotyping as a scalable tool for glioma prognostication.

The central methodological innovation of this work is the transfer of molecularly resolved cell type annotations from Xenium spatial transcriptomics data to matched H&E images, enabling supervision of a detection model with molecular ground truth rather than morphologic heuristics alone. This cross-modal strategy, previously demonstrated in lung cancer brain metastases and other tumor contexts^14,15^, is particularly well-suited to vascular biology, where the distinction between endothelial cells, pericytes, and perivascular fibroblasts is biologically meaningful but morphologically ambiguous on routine staining. By encoding this molecular specificity into an H&E-based model, our approach bridges the resolution gap between resource-intensive spatial omics and the practical scalability demanded by large retrospective archives. Importantly, we demonstrate that annotations derived from a small set of WSIs can be sufficient to train a performant detection model when spatial transcriptomics provides high-fidelity, cell-level ground truth, circumventing the need for large manually annotated training sets that are impractical to generate for rare cell populations such as vascular cells.

The binary vascular detection model achieved a precision of 0.78 and an F1 score of 0.70, with an overall accuracy of 98.6%. The recall of 0.63, while modest, reflects a well-characterized challenge in rare-cell detection: vascular cells constitute approximately 1.2% of all annotated cells in the training tissue, creating substantial class imbalance that biases the model toward the majority class. Orthogonal spatial validation in four independent Xenium patient slides confirmed that the model’s detections were biologically coherent, with predicted vascular cells showing significantly greater proximity to Xenium-annotated endothelial and pericyte cells than expected by chance under a permutation-based null distribution. This spatial fidelity is critical for translational credibility and distinguishes our approach from models trained on morphologic surrogates alone.

The most clinically significant finding of this study is the independent prognostic value of AI-derived vascular proportion in astrocytoma. Astrocytomas span WHO grades 2 and 3 and represent the glioma subtype where prognostic stratification beyond grade is most clinically consequential, as grade alone incompletely predicts outcome heterogeneity within this group^23,3^. The 2.3-fold increased hazard associated with high vascular proportion, maintained after adjustment for age and sex, suggests that the angiogenic phenotype captured by our model reflects a biologically aggressive subtype of astrocytoma that is not fully resolved by current histologic classification. This is consistent with the known role of tumor vasculature in sustaining glioma stem cell niches and driving progression independent of grade^4,6^ .

Several limitations of this study merit acknowledgement. First, the detection model was trained exclusively on a single glioblastoma Xenium specimen, which may limit representation of the full spectrum of vascular morphologies encountered across glioma subtypes and grades. In particular, the lower-grade vascular phenotypes characteristic of astrocytoma and oligodendroglioma, including delicate chicken-wire vasculature and early angiogenic changes distinct from the florid microvascular proliferation of GBM, were not represented in the training data. Expanding the training set to include Xenium data from lower-grade tumors is likely to improve model sensitivity and morphologic specificity across the glioma spectrum. Second, the orthogonal validation dataset comprised four patient slides from a single published cohort, which represents a limited external validation set. Broader multi-institutional validation across diverse tissue preparation and scanning protocols is necessary to establish the robustness of the model before clinical deployment. Third, the retrospective cohort of 119 cases, while sufficient for hypothesis generation, is modest in size for survival analysis, particularly within subtype strata; the oligodendroglioma group (n = 21) was too small to draw meaningful conclusions. Prospective validation in larger, independently collected cohorts is required before clinical translation. Finally, the current model classifies cells as vascular or non-vascular but does not distinguish between histologically and prognostically distinct vascular subtypes, such as chicken-wire vasculature, glomeruloid microvascular proliferation, or endothelial hypertrophy, which carry different diagnostic and prognostic implications under current WHO criteria^2^. Future work extending the classification framework to resolve these vascular subtypes would substantially increase the clinical utility of the approach.

Despite these limitations, this study establishes a methodologically rigorous framework for molecularly informed, spatially supervised vascular phenotyping at the histologic scale. The approach is inherently scalable; once trained, the model can be applied to any H&E WSI without additional molecular profiling, and is compatible with existing clinical pathology workflows. As spatial transcriptomics datasets continue to grow in scope and diversity, cross-modal training strategies of this kind offer an increasingly powerful means of encoding molecular tissue biology into universally available histologic material, ultimately enabling prognostic and predictive biomarkers to be extracted from the vast archives of routine pathology that would otherwise remain beyond the reach of molecular analysis.

## Author Contributions

Dr. Nameeta Shah and Dr. Megha S. Uppin conceived and designed the study. Ramya Alugam performed data collection, slide review, and annotation. Pranali S and Dr. Nameeta Shah conducted data analysis. Shashank Gupta contributed to AI model development. Pranali S drafted the manuscript. Dr. Megha S. Uppin and Dr. Nameeta Shah critically reviewed and edited the manuscript. All authors reviewed and approved the final version of the manuscript.

## Ethics Approval and Consent

Ethical approval for this study was obtained from the Institutional Ethics Committee of Nizam’s Institute of Medical Sciences (NIMS-IEC) (Reference No: EC/NIMS/3259/2023). All patient-related information, including pathology case identification numbers, was anonymized, and new case numbers were assigned to ensure confidentiality and protection of patient rights in accordance with institutional ethical guidelines.

## Acknowledgements

We acknowledge the Indian Council of Medical Research (ICMR) for providing financial assistance to support this research work.

## Code Availability

No custom code was developed for this study. Model training was performed using the Amaranth Insights no-code AI platform (https://www.amaranth-insights.com/), which implements a DINO-DETR-based object detection architecture. Survival analyses were conducted in R (version 4.3.1) using the survival and survminer packages.

## Data Availability

The 10x Xenium spatial transcriptomics image used for model training is a publicly available demonstration dataset provided by 10x Genomics (https://www.10xgenomics.com/datasets). The Xenium slides used for external model validation were derived from the glioblastoma atlas published by Sonpatki et al. (Nat Commun, 2026; https://doi.org/10.1038/s41467-026-69716-2); the associated data are publicly available at https://doi.org/10.5281/zenodo.17622242. The H&E WSIs from the retrospective cohort and AI-derived vascular annotations are accessible at https://www.amaranth-insights.com/. Clinical and outcome data are available from the corresponding author on reasonable request, subject to institutional ethics approval.

## References

1. Loureiro LVM, Neder L, Callegaro-Filho D, et al. The immunohistochemical landscape of VEGF family and its receptors in glioblastomas. Surg Exp Pathol. 2020;3:9.

2. Louis DN, Perry A, Wesseling P, et al. The 2021 WHO Classification of Tumors of the Central Nervous System: a summary. Neuro Oncol. 2021;23(8):1231–1251.

3. Jain R, Poisson L, Narang J, et al. Genomic mapping and survival prediction in glioblastoma. Radiology. 2013;267(1):173–183.

4. Calabrese C, Poppleton H, Kocak M, et al. A perivascular niche for brain tumor stem cells. Cancer Cell. 2007;11(1):69–82.

5. Friedman HS, Prados MD, Wen PY, et al. Bevacizumab alone and in combination with irinotecan in recurrent glioblastoma. J Clin Oncol. 2009;27(28):4733–4740.

6. Jhaveri N, Chen TC, Hofman FM. Tumor vasculature and glioma stem cells: contributions to glioma progression. Cancer Lett. 2016;380(2):545–551.

7. Fox SB, Harris AL. Histological quantitation of tumour angiogenesis. APMIS. 2004;112(7–8):413–430.

8. Cheng J, Jin X, Smyth GK, Chen Y. Benchmarking cell type annotation methods for 10x Xenium spatial transcriptomics data. BMC Bioinformatics. 2025;26(1):22.

9. Bressan D, Battistoni G, Hannon GJ. The dawn of spatial omics. Science. 2023;381(6657):eabq4964.

10. Williams CG, Lee HJ, Asatsuma T, Vento-Tormo R, Haque A. An introduction to spatial transcriptomics for biomedical research. Genome Med. 2022;14(1):68.

11. Rao A, Barkley D, França GS, Yanai I. Exploring tissue architecture using spatial transcriptomics. Nature. 2021;596(7871):211–220.

12. Chen X, Lin J, Wang Y, et al. HE2Gene: image-to-RNA translation via multi-task learning for spatial transcriptomics data. Bioinformatics. 2024;40(6):btae343.

13. Janesick A, Shelansky R, Gottscho AD, et al. High resolution mapping of the tumor microenvironment using integrated single-cell, spatial and in situ analysis. Nat Commun. 2023;14(1):8353.

14. Danaher P, Kim Y, Nelson B, et al. Jointly leveraging spatial transcriptomics and deep learning models for pathology image annotation improves cell type identification over either approach alone. bioRxiv. 2021. doi:10.1101/2021.11.10.468082.

15. Gavish A, Baron M, Chomsky E, et al. The spatial transcriptomic landscape of non-small-cell lung cancer brain metastases. bioRxiv. 2023.

16. Carion N, Massa F, Synnaeve G, et al. End-to-end object detection with transformers. ECCV. 2020;12346:213–229.

17. 10x Genomics. FFPE Human Brain Cancer Data with Human Immuno-Oncology Profiling Panel and Custom Add-on. Xenium Onboard Analysis v2.0.0. Published April 15, 2024. Available at: https://www.10xgenomics.com/datasets/ffpe-human-brain-cancer-data-with-human-immuno-oncology-profiling-panel-and-custom-add-on-1-standard.

18. Redlich JP, Feuerhake F, Weis J, et al. Applications of artificial intelligence in the analysis of histopathology images of gliomas: a review. npj Imaging. 2024;2(1):16.

19. Chunduru P, Phillips JJ, Molinaro AM. Prognostic risk stratification of gliomas using deep learning in digital pathology images. Neurooncol Adv. 2022;4(1):vdac111.

20. Liu Z, Li X, Zhao F, et al. A multi-center performance assessment for automated histopathological classification and grading of glioma using whole slide images. iScience. 2023;26(11):108196.

21. Zhang H, Li F, Liu S, et al. DINO: DETR with improved denoising anchor boxes for end-to-end object detection. ICLR. 2023.

22. Sonpatki P, Park HJ, Xing YL, et al. A spatially resolved human glioblastoma atlas reveals distinct cellular and molecular patterns of anatomical niches. Nat Commun. 2026. doi:10.1038/s41467-026-69716-2.

23. Cheng J, Liu X, et al. The spectrum of microvascular patterns in adult diffuse glioma and their correlation with tumor grade. J Pathol Transl Med. 2024;58:108–117.

24. Chauhan E, Sharma A, Uppin MS, Kondamadugu M, Jawahar CV, Vinod PK. IPD-Brain: An Indian histopathology dataset for glioma subtype classification. Scientific Data. 2024;11:1403. 10.1038/s41597-024-04225-9.

25. Jha K, Pant I, Singh R, Bansal AK, Chaturvedi S. Assessment of microvascular patterns and density in glioblastoma and their correlation with prognosis. J Neurooncol. 2014.

26. Tena-Suck ML, Salinas-Lara C, Arce-Arellano RI, et al. Glioblastoma multiforme and angiogenesis: a clinicopathological and immunohistochemistry approach. J Neurol Res. 2015;5(3):166–176.

